# SLAM-seq reveals independent contributions of RNA processing and stability to gene expression in African trypanosomes

**DOI:** 10.1101/2024.06.18.599538

**Authors:** Vanessa Luzak, Esteban Osses, Anna Danese, Christoff Odendaal, Stefan H. Stricker, Jurgen R. Haanstra, Florian Erhard, T. Nicolai Siegel

**Affiliations:** Division of Experimental Parasitology, Faculty of Veterinary Medicine, Ludwig-Maximilians-Universität München, Munich, Germany; Biomedical Center, Division of Physiological Chemistry, Faculty of Medicine, Ludwig-Maximilians-Universität München, Munich, Germany; Reprogramming and Regeneration, Biomedical Center (BMC), Physiological Genomics, Faculty of Medicine, Ludwig Maximilian University (LMU) Munich, Planegg-Martinsried 82152, Germany; Epigenetic Engineering, Institute of Stem Cell Research, Helmholtz Zentrum, German Research Center for Environmental Health, Planegg-Martinsried 82152, Germany; Systems Biology Lab/A-LIFE, Amsterdam Institute of Molecular and Life Sciences (AIMMS), Vrije Universiteit Amsterdam, Amsterdam, The Netherlands; Institut für Virologie und Immunbiologie, Julius-Maximilians-Universität Würzburg, Würzburg, Germany; Chair of Computational Immunology, University of Regensburg, 93053 Regensburg, Germany

## Abstract

Gene expression is a multi-step process that converts DNA-encoded information into proteins, involving RNA transcription, maturation, degradation, and translation. While transcriptional control is a major regulator of protein levels, the role of post-transcriptional processes such as RNA processing and degradation is less well understood due to the challenge of measuring their contributions individually.

To address this challenge, we investigated the control of gene expression in *Trypanosoma brucei*, a unicellular parasite assumed to lack transcriptional control. Instead, mRNA levels in T*. brucei* are controlled by post-transcriptional processes, which enabled us to disentangle the contribution of both processes to total mRNA levels.

In this study, we developed an efficient metabolic RNA labeling approach and combined ultra-short metabolic labeling with transient transcriptome sequencing (TT-seq) to confirm the long-standing assumption that RNA polymerase II transcription is unregulated in *T. brucei*. In addition, we established thiol (SH)-linked alkylation for metabolic sequencing of RNA (SLAM-seq) to globally quantify RNA processing rates and half-lives. Our data, combined with scRNA-seq data, indicate that RNA processing and stability independently affect total mRNA levels and contribute to the variability seen between individual cells in African trypanosomes.

## Introduction

Gene expression is a multi-step process (1) that converts the information encoded by a gene into messenger RNA (mRNA) and subsequently into proteins. Eukaryotic genomes encode thousands of proteins, each of which is required at different levels in the cell. While some proteins, such as chromatin-modifying enzymes, are present at low levels of ∼100 copies per cell, others, such as ribosomal subunits, are present at 1,000-fold higher levels (2). To achieve appropriate levels of each protein, regulation can occur at each step of the gene expression cascade: (1) The rate of transcription determines how often a gene is transcribed into pre-RNA (3, 4). (2) The efficiency of co-transcriptional mRNA processing by splicing and polyadenylation determines what fraction of pre-RNA is converted into mature mRNA and what fraction is prematurely degraded (5–7). (3) The stability of mature mRNAs then determines how rapidly mRNAs are degraded (8, 9). Together, transcription, RNA processing and RNA stability control the pool of RNAs in a cell. In addition, control of gene expression also occurs during protein synthesis: (4) Translation efficiency refers to the rate at which an mRNA is translated into protein and varies widely among mRNAs(10, 11). (5) The rate of protein processing, such as post-translational modifications or proteolytic processes, can control the amount of functionally active protein in a cell (12, 13). (6) Finally, protein stability determines the rate at which a given protein is degraded (14, 15).

A common strategy for assessing gene expression is to measure total mRNA levels, typically using high-throughput RNA-sequencing (RNA-seq) (16). While RNA-seq is a powerful approach to accurately determine total mRNA levels on a genome-wide scale, it does not provide information on how transcription, RNA processing, and stability contribute to the measured RNA levels. However, in the physiological context of a cell, the step at which the expression of a transcript is controlled makes a difference. Control at the transcriptional level might be energy efficient, but volatile and slow (17). In most eukaryotes, transcription of individual genes is controlled by a gene-specific promoter sequence that recruits transcription factors and can be further fine-tuned by cis-regulatory enhancer sequences(18). In contrast, controlling gene expression at the level of RNA stability allows the cell to confer robustness to the expression of a gene. A long mRNA half-life can buffer the intrinsic noise of other sub-steps in the gene expression cascade, while a short mRNA half-life allows a cell to quickly adapt to environmental changes. The stability of mature mRNA is controlled in several ways, including by RNA-binding proteins that bind sequence motifs in the untranslated regions (UTRs) of mRNAs, thereby affecting RNA transport, localization, and storage (19–21). In addition to sequence motifs in the UTRs, codon usage by the protein coding sequence (CDS) of an mRNA influences its stability by controlling translation efficiency and ribosome occupancy of a transcript (22–24).

While the control of gene expression at the level of transcription and RNA stability has long been appreciated and is well understood in different organisms, it is less well understood how gene-specific RNA processing rates control total mRNA levels. Only recently, differences in processing efficiency have been systematically measured in mammalian cells and were suggested as an additional player of control (6). However, it has remained unclear how the two post-transcriptional processes, RNA processing and RNA stability, ‘interact’ and whether they cooperate or antagonize to regulate total mRNA levels for different transcripts. Assuming regulatory independence per transcript, there should be two sets of transcripts with intermediate total levels: one set of transcripts processed at high rates, but with low RNA stability, and another set of transcripts with low processing rates and high stability. Importantly, each set of transcripts should be able to play different functional roles. Genes with high RNA stability are robustly expressed in all cells, while genes with high RNA processing rates and low RNA stability could enable the cell to rapidly adapt to environmental changes.

To investigate the combined impact of RNA processing rates and RNA stability on total mRNA levels, we used *Trypanosoma brucei* as a eukaryotic model in which transcriptional control is thought to be absent. *T. brucei* is a highly divergent, unicellular parasite responsible for sleeping sickness in humans and nagana in cattle in sub-Saharan Africa. Unusually for a eukaryote, its ∼9,000 genes transcribed by RNA polymerase II are organized into ∼200 polycistronic transcription units (PTUs) (25, 26), and PTU transcription is thought to occur without gene-specific regulation in an unregulated, constitutive manner (4). Furthermore, since introns have only been reported for two genes (27), there are no complex cis-splicing patterns in *T. brucei*. Instead, pre-mRNA processing in *T. brucei* involves polyadenylation and a *trans*-splicing step that adds a common spliced leader sequence to each pre-mRNA (7, 28). Taking advantage of the apparent lack of transcriptional control and a less complex RNA processing mechanism, we used *T. brucei* to disentangle the effects of RNA processing rates and RNA stability on total mRNA levels.

In this study, we have established an efficient approach for the metabolic labeling of newly synthesized RNA in bloodstream form trypanosomes. Efficient labeling is achieved by genetic manipulation of the endogenous pyrimidine synthesis pathway. By integrating ultra-short metabolic labeling with transient transcriptome sequencing (TT-seq) (29, 30) we were able to corroborate the long-standing assumption that RNA polymerase II transcription is a rather unregulated process in *T. brucei*. Furthermore, we established thiol(SH)-linked alkylation for the metabolic sequencing of RNA (SLAM-seq) (31, 32) to assess RNA processing rates and RNA half-lives on a genome-wide scale. Our results indicate that post-transcriptional processes operate in an independent manner and play an important role in cell-to-cell variability in African trypanosomes.

## Materials and methods

### Cell culture

*Trypanosoma brucei* cells derived from the Lister 427 bloodstream form strain MiTat 1.2 (clone 221a) strain were cultured in HMI-11 medium (HMI-9 medium(33) without serum-plus) at 37 °C and 5% CO_2_. The following drug concentrations were used when appropriate: 2 µg ml^−1^ G418 (neomycin), 5 µg ml^−1^ hygromycin, 0.1 μg ml^−1^ puromycin, 5 μg ml^−1^ blasticidin, 2.5 µg ml^−1^ phleomycin and 1 μg ml^−1^ doxycycline (DOX).

### Cell line generation

Transfections were performed using a Nucleofector device (Amaxa) and following a protocol established for *T. brucei* (34). The UMPS KO cell line was generated from “single marker” (SM) cells, expressing T7 polymerase and a Tet repressor (35), by replacing both UMPS alleles with resistance markers. Two previously published plasmids (34) were used for UMPS replacement: pyrFEKO-HYG and pyrFEKO-PUR. Subsequently, the resistance markers were removed by transient Cre recombinase expression (34).

The FCU overexpression (36) construct was transfected into the “2T1 T7” strain, which expresses a Tet repressor, T7 polymerase, and contains a tagged ribosomal spacer as a landing pad for the transfection construct (37), and into the UMPS KO cell line. For FCU expression, the plasmids pVLFCU (targeting the 2T1 T7 landing pad; no 3xHA-tag) and pVL_FCU_2 (targeting a random rDNA spacer; 3x HA-tag) were generated, using the previously published plasmids pRPCas9 and pT7sgRNA as parental plasmids, respectively (38). pRPCas9 was digested with HindIII and XhoI, and pT7sgRNA was digested with NheI and XbaI. The FCU coding sequence (CDS) was ordered as gBlock GB37_FCU_CDS from IDT and inserted into the digested pRPCas9 plasmid by infusion cloning. For insertion into the digested pT7sgRNA plasmid, the FCU CDS was amplified from the pVLFCU plasmid using the FCU_for and FUR1_CDS_rev primers. In addition, a fragment carrying the 5’UTR was amplified from pVLFCU using the primers FUR1_UTR_for and FUR1_UTR_rev. Both PCR products, carrying the FCU CDS and the 5’UTR, were joined to the digested pT7sgRNA backbone by infusion cloning, as well as a short fragment to add an N-terminal 3x HA-tag.

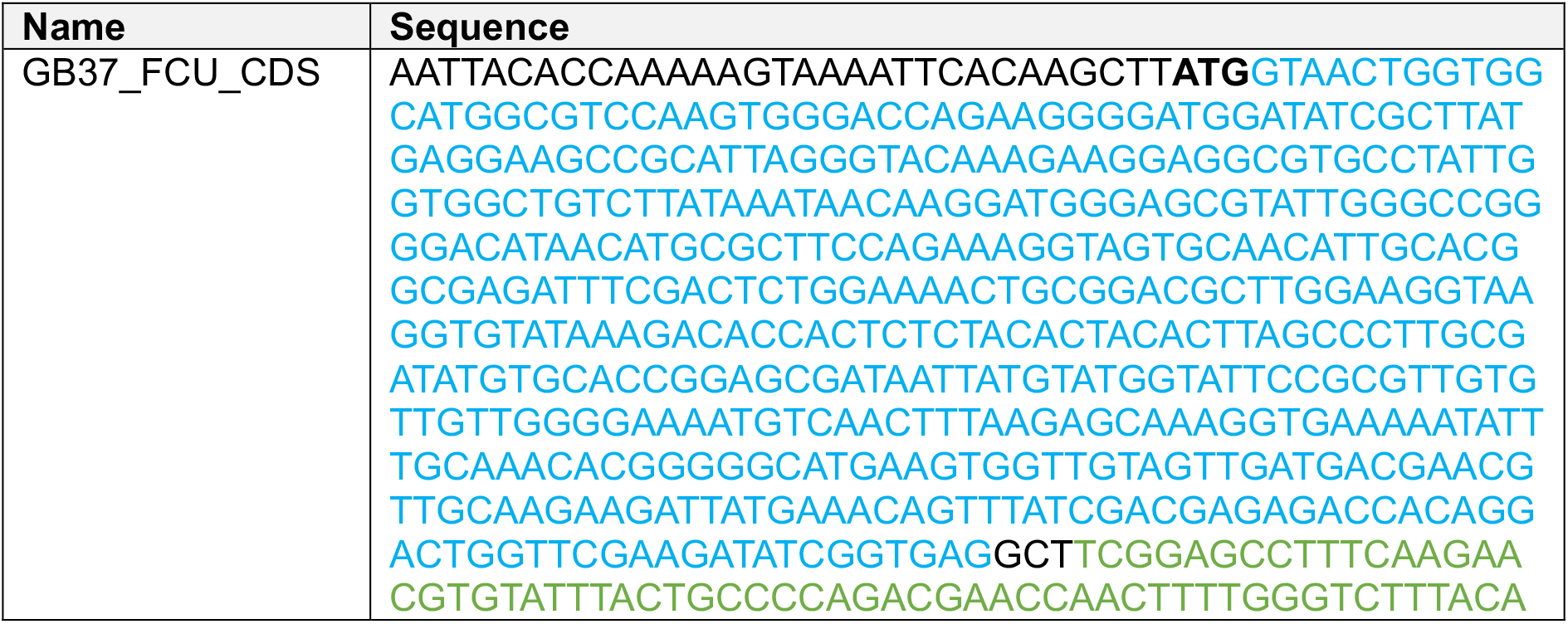
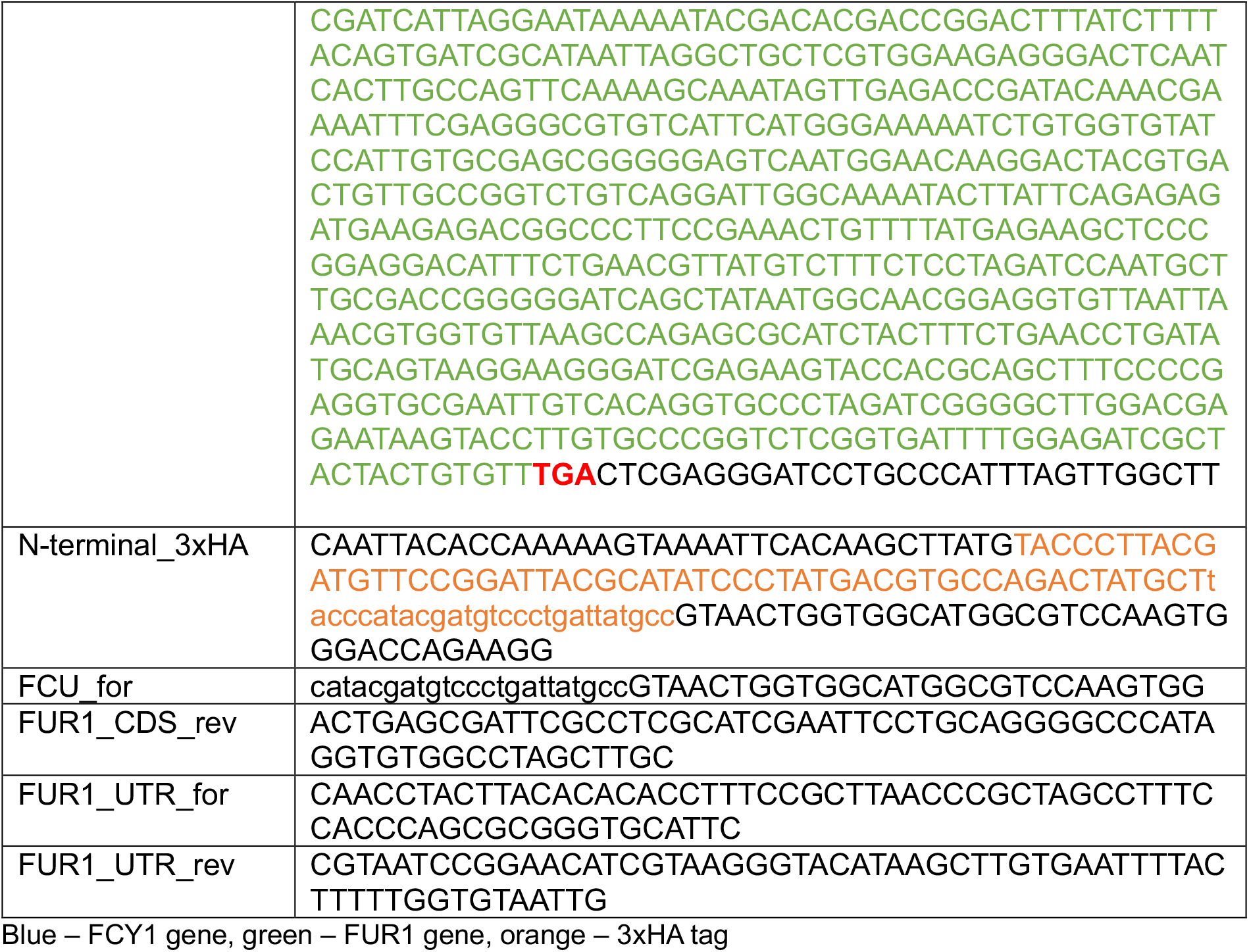

### Western blot

Expression of the 3xHA-tagged FCU protein was induced by doxycyclin for 24 hours and verified by western blot analysis in three independent clones. *T. brucei* cells were harvested by centrifugation for 5 minutes at 1000 x g and lysed in RIPA buffer (50 mM Tris-HCl pH 8.0, 150 mM NaCl, 1 % NP-40, 0.25% sodium-deoxycholate, 0.1 % SDS) supplemented 1:3 with 4x Laemmli buffer (40% glycerol 0.02 % bromphenol blue, 8 % SDS, 250 mM Tris pH 6.5). Extracts equivalent to 2 × 10^6^ cells were separated on 10 % SDS-polyacrylamide gels and transferred to a nitrocellulose membrane (Amersham 10600001) by wet transfer. Total protein was visualized with Amidoblack 10b staining solution to ensure even loading and protein transfer. The blots were blocked in 3% BSA in PBS-T and washed three times in PBS-T (0.05% Tween). Next, the blots were cut in two, and the upper part (above 25 KDa according to the pre-stained protein ladder, Thermo Fisher 26619) was probed with 1/1000 α-HA primary antibody (Sigma H6908) to detect 3xHA_FCU, and 1/10000 α-rabbit secondary HRP antibody (Thermo Fisher, NA934V). As a loading control, the bottom part (below 25 KDa according to the pre-stained protein ladder) was incubated with 1/1000 α-H2A.Z (39) and 1/10000 α-rabbit secondary HRP antibody (Thermo Fisher, NA934V).

### Metabolic RNA labeling and ultra-fast RNA harvest

Prior to metabolic labeling of newly synthesized RNA, bloodstream form cells were grown to a density of 0.6 - 0.8 million cells/ml in standard HMI-11 medium, in order to ensure an exponential growth rate during the labeling period. For metabolic labeling with 4-TU (4-thiouracil, Sigma Aldrich, 440736), the cell suspension was transferred to 50 ml Falcons for labeling in a water bath at 37 °C. If cells were labeled in a pyrimidine-reduced version of the HMI-11 medium (without thymidine and with dialyzed FCS (40)), a centrifugation step was added at this point for 5 minutes at 1000 x g. After pelleting the cells, the standard medium was discarded and replaced with 50 ml of pre-warmed pyrimidine-reduced medium. The required amount of a 200 mM 4-TU stock solution in DMSO was added to the cell suspension in the Falcon and mixed by inversion. The Falcon was placed in the water bath and labeling was performed at 37 °C for the intended labeling time. At the end of the labeling period, the cell suspension was immediately passed through a 0.8 μm hydrophilic membrane (MF-Millipore, AAWP04700) mounted on a Pyrex suction filter (PORO3 5810/3 40/38, 5809/2, 40/38 5810/4) connected to vacuum. The membrane was then quickly transferred to a fresh 50 ml Falcon and placed in liquid nitrogen, to ensure ultra-short harvest times. Membranes were transferred to -80 °C until further processing. For RNA harvest, the membranes were removed from -80 °C and thawed for 1 minute at RT. Two ml Trizol reagent (Thermo Fisher 15596-026) was added per membrane and cell lysis was completed by incubation on a tube roller for 10 minutes. Cell lysates in Trizol were transferred to 2 ml Eppendorf tubes and a standard isopropanol extraction of RNA was performed.

### Dot blot

For dot blot analysis, 3 µg of total mRNA was biotinylated per sample and three rounds of chloroform washes were performed to remove free biotin, as previously described (41). 1,200 ng and 300 ng of RNA were spotted onto a Hybond N+ Amersham membrane (RPN203B) and UV cross-linked at 120 mJ. The membrane was incubated in blocking buffer (phosphate buffered saline (1xPBS), pH 7.5, 10% SDS, 1 mM EDTA) for 20 minutes. After blocking, a 1:1000 dilution of 1 mg/ml streptavidin-HRP antibody (Abcam 7403) was added to the blocking buffer for 15 minutes. Six washes were then performed: 2x with 10% SDS in PBS, 2x with 1% SDS in PBS and 2x with 0.1% SDS in PBS.

### SLAM-seq: chemical conversion and library preparation

SLAM-seq analysis was performed to quantify the efficiency of the different labeling strategies tested in Figure 1. Forty million cells were used per sample and labeling was performed for three hours to reach saturation, in either standard HMI-11 medium or a pyrimidine-reduced version (40) (no thymidine added and dialyzed FCS). Further, SLAM-seq analysis was performed to determine RNA half-lives and RNA synthesis rates in bloodstream parasites, using the following conditions: 40 million cells were used per replicate and three biological replicates were prepared for each time point. Labeling with 4-TU was performed for 0, 7.5, 15 and 60 minutes in standard HMI-11 medium, in order to capture the full range of RNA half-lives estimated from a previous experiment (9). Metabolic labeling and RNA harvest were performed as described above.

**Figure 1.**
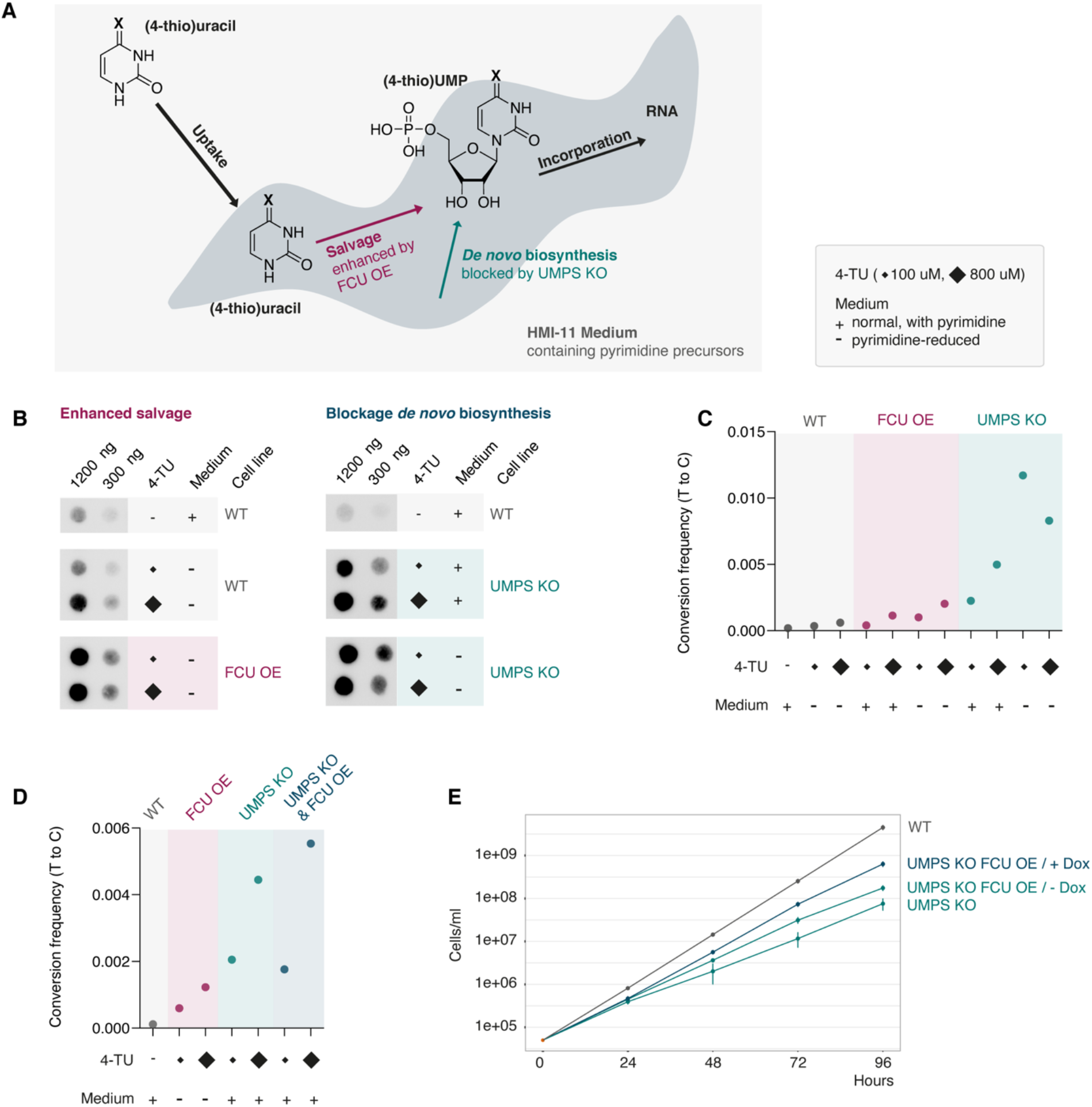
Genetic manipulation of pyrimidine metabolism in *T. brucei* enables efficient metabolic RNA labeling. **(A)** Schematic illustration of the two main pathways of pyrimidine synthesis in *T. brucei*: salvage of external precursors and *de novo* biosynthesis. **(B)** Dot blot analysis to detect 4-TU incorporation into *T. brucei* RNA was performed in wild-type cells (WT), after enhancement of the salvage pathway by FCU overexpression (OE) and after blocking *de novo* biosynthesis by UMPS knockout (KO). Incubation time 3 hours. **(C)** SLAM-seq analysis was performed to quantify 4-TU incorporation efficiency by measuring T-to-C conversion frequencies in WT, FCU OE, and UMPS KO cells. Incubation time 3 hours. **(D)** SLAM-seq analysis to evaluate the effect of the combination of FCU OE and UMPS KO on pyrimidine incorporation. **(E)** Growth curves of WT, UMPS KO, UMPS KO cells expressing FCU. Growth curves were performed in triplicate and error bars indicate the standard deviation at each time point. FCU expression was induced by doxycycline (Dox) for 24 hours prior to growth rate analyses.

Chemical conversion was performed according to Herzog *et al*. (32), with minor modifications: the conversion reaction was performed for 30 minutes instead of 15 minutes, in order to obtain uniform conversion rates. rRNA depletion was performed according to an RNA-seq protocol previously established for *T. brucei* by the Siegel group (42). Library preparation was performed using the NEBNext® Ultra II Directional RNA Library Prep Kit for Illumina (NEB #E7760S/L). The workflow was adapted to generate longer library fragments by reducing the fragmentation time (see manual, Appendix A, section 6.1) to increase the accuracy of SNP detection. 30 million reads were sequenced per replicate at 150 bps paired end.

Data analysis to determine RNA half-lives and synthesis rates was performed using the GRAND-SLAM (43) and grandR (44). In brief: reads were mapped to the *T. brucei* Lister 427 2018 genome assembly (release 36, downloaded from TritrypDB) using STAR (45) (version 2.5.3a) using parameters “–outFilterMismatchNmax 40–outFilterScoreMinOverLread 0.2– outFilterMatchNminOverLread 0.2–alignEndsType Extend5pOfReads12–outSAMattributes nM MD NH”. Bam files were merged and converted into a CIT file using the GEDI toolkit (46) and then processed using GRAND-SLAM 2.0.7 (43) to generate read counts and NTR values on the gene level. The grandR (44) tool was used to estimate RNA half-lives (minutes) and synthesis rates (TPM/hour) using the non-linear least squares approach. Assuming unregulated transcription in trypanosomes, the synthesis rate was used to describe mRNA processing rates. The synthesis rate was initially calculated in TPM/hour, and converted into molecules/hour assuming 33,300 transcripts per cell. This was calculated using the estimate from Fadda *et al.*(9) of 20,000 mRNA molecules.cell^-1^ corrected for the fact that in the SLAM-seq experiment 60% of transcripts were mRNA. Further, the GO-term analysis for the four gene groups resulting from RNA half-life and processing regulation was performed with TritrypDB (47) and visualization was performed using the GO-figure (48) tool.

### TT-seq: biotinylation, enrichment and library preparation

For TT-seq analysis, 700 million cells per replicate were used and the experiment was performed in duplicate. Ultra-short labeling with 4-TU was performed for 0, 30, 60 and 120 seconds, in order to capture newly synthesized RNA (49). Labeling was performed in a pyrimidine-reduced version of the HMI-11 medium as described above. Subsequent biotinylation and enrichment using streptavidin beads were performed as previously described (50). rRNA depletion was performed on input samples, but not on IP samples due to the low amount of material in nascent RNA samples. Library preparation was performed using the NEBNext® Ultra II Directional RNA Library Prep Kit for Illumina (NEB #E7760S/L) as described for standard RNA-seq (42). Thirty million reads were sequenced per replicate at 60 bps paired end.

After quality control with FastQC, reads were mapped to the *T. brucei* Lister 427 2018 genome assembly (release 36, downloaded from TritrypDB) using BWA-mem (51). The resulting sam files were converted to bam format and sorted and indexed using SAMtools 1.8 (52). Additionally, unmapped, PCR or optical duplicates, non-primary aligned and supplementary aligned reads were filtered out from the alignment files (SAM flag: 3332). For visualization, bigwig, bedgraph and metaplot files were generated from the sorted bam files using deepTools 3.5.0 (53). Reads mapped to the genome were plotted using PyGenome-track (54). Read count tables were generated using Subread 1.6.2 (55) and PCA analysis was performed using the Bioconductor package DESeq2 (56).

### RNA-seq

RNA-seq analysis was performed as previously described for *T. brucei*, using specific biotinylated oligos for rRNA removal (42). Ten million reads were sequenced per replicate at 60 bps paired end. After quality control with FastQC, reads were mapped to the *T. brucei* Lister 427 2018 genome assembly (release 36, downloaded from TritrypDB) using BWA-mem (51). The resulting sam files were converted to bam format and sorted and indexed using SAMtools 1.8 (52). In addition, unmapped, PCR or optical duplicate, non-primary aligned and supplementary aligned reads were filtered out of the alignment files (SAM flag: 3332). Read count tables were generated using Subread 1.6.2 (55). PCA and differential expression analysis were performed using the Bioconductor package DESeq2 (56).

### Modeling analysis

A computational model of gene expression in the bloodstream form of *T. brucei* was adapted from Antwi *et al.* (57). The model was built in Python (version 3.10.9) using the kinetic modelling package PySCeS (58) (version 1.1.1). The model was reduced to three reactions: an mRNA synthesis reaction (v_synth_), an mRNA degradation reaction (v_degr_) and growth-rate dilution (v_dil_). v_synth_ replaces the transcription and processing reactions in the previous model and has a constant value, while mRNA concentration-dependent mass-action kinetics is used for the other two reactions:

> v_degr_ = k_degr_*[mRNA]

> v_dil_ = µ*[mRNA]

All rates pertain only to mRNA levels and have units of mRNA molecules.cell^-1^.min^-1^, just as mRNA concentration has units of molecules/cell. The specific growth rate (µ) was set to 0.0019 min^-1^, as measured by Haanstra *et al.* (59). The default value for v_synth_ was set to 0.1667 mRNA molecules.min^-1^, which is the simulated mRNA production flux in Antwi *et al.*’s (57) bloodstream form A model. The default k_degr_ was 0.015 min^-1^ as calculated for PGKC mRNA by Haanstra *et al.* (59). For the transcript-specific models, the values for v_synth_ and k_degr_ were replaced by the relevant newly measured values. For the conversion from molecules/cell to TPM, we assumed 33,300 transcripts per cell. This was calculated using the estimate from Fadda *et al.* (9) of 20,000 mRNA molecules.cell^-1^ corrected for the fact that in the SLAM-seq experiment 60% of transcripts were mRNA.

### Single-cell RNA-seq analysis

The single cell RNA-seq data from bloodstream form trypanosomes was generated previously (60). For this study, reads were mapped to an allele-specific, phased genome version (25) using zUMIs (61) (version 2.9.7) with STAR (45) (version 2.7.10). Cells were filtered out when less than 150 genes were detected, and genes were filtered out when detected in less than 10 cells, resulting in 369 cells and 7,163 detected genes for the scRNA-seq dataset. Of the 7,163 genes detected by scRNA-seq, the previous SLAM-seq experiment had revealed RNA half-life and processing rate for 6,176 genes, which were used for further analysis. To estimate the variation of gene expression in individual cells, the mapped single cell RNA-seq data was normalized to RPKM (reads per kilobase of transcript per million reads mapped) using Scanpy (62). Genes were divided into four groups according to their RNA stability and RNA processing rates, as described above. Then, the cell-to-cell variability of normalized gene expression was determined by calculating the coefficient of variation across cells. Further, for all genes, both, bi-allelic and allele-specific counts were generated, and a generalized telegraph model (GTM) was run, first described by Luo *et al.* (63). Using a python re-implementation of the GTM (https://github.com/DaneseAnna/transcriptional_bursting_dev), we obtained bi-allelic and mono-allelic transcript expression parameters.

## Results

### Genetic manipulation of pyrimidine metabolism enables efficient RNA labeling

To gain insight into the gene expression process beyond that provided by conventional RNA-seq assays, we first established an approach for efficient metabolic labeling of newly synthesized RNA in bloodstream form *T. brucei*, responsible for infections in mammals. A sufficiently high incorporation rate of nucleotide analogues during metabolic RNA labeling is required to reliably differentiate between new and old RNA and to allow a detailed analysis of the different gene expression steps. While the less versatile uridine analog 5-EU has been incorporated at detectable rates in insect-stage trypanosomes (64), incorporation of uridine analogues to detectable levels has not been possible in bloodstream form parasites (65). A possible reason may be the less efficient uridine uptake in this life cycle stage (40). We therefore chose the uracil analog 4-TU for our experiments, rather than the uridine-based agents 4-SU or 5-EU, because uracil has been reported to be more efficiently taken up from the medium by bloodstream form cells than uridine (40).

Furthermore, we used a two-pronged genetic approach to increase the incorporation of pyrimidine analogues into nascent RNA (Figure 1A): On the one hand, to enhance the intracellular salvage pathway of pyrimidine precursors from the medium, we exogenously expressed the yeast cytosine deaminase-uracil phosphoribosyl transferase fusion enzyme FCU (Supplementary Figure S1A). This enzyme converts 4-thiouracil to 4-thioUMP and has been used, for example, in *Plasmodium* parasites to increase the efficiency of metabolic RNA labeling (36) . On the other hand, to block *de novo* biosynthesis of pyrimidines, we deleted both alleles of the endogenous uridine monophosphate synthase (UMPS) gene (Supplementary Figure S1B). UMPS encodes an enzyme that is critical for the *de novo* biosynthesis pathway, and the knockout results in a complete block of *de novo* biosynthesis. However, it does not interfere with the salvage pathway (40, 66).

Metabolic labeling was performed with 100 µM and 800 µM 4-TU in a) wild-type bloodstream form cells, b) cells overexpressing the FCU enzyme (enhanced pyrimidine salvage), and c) UMPS knockout cells (blocked *de novo* pyrimidine biosynthesis). Labeling was performed for three hours to reach saturation, both in normal HMI-11 medium containing pyrimidine precursors and a pyrimidine-reduced version of HMI-11 (40). Subsequently, total cellular RNA was extracted and 4-TU incorporation was assessed by biotinylation and dot blot analysis (Figure 1B), as well as by chemical conversion and Illumina sequencing (Figure 1C). Based on dot blot signal intensity, both strategies, enhancing pyrimidine salvage or blocking pyrimidine *de novo* biosynthesis, increased nascent RNA labeling compared to wild-type cells. Quantification by SLAM-seq showed that blocking pyrimidine *de novo* biosynthesis was more efficient than enhancing pyrimidine salvage (Figure 1C). As expected, labeling was more efficient in pyrimidine-reduced medium than in standard medium, and for most conditions, incorporation rates were higher when labeling with 800 µM 4-TU compared to 100 µM 4-TU, except for one condition. When UMPS knockout cells were used for labeling in pyrimidine- reduced conditions, labeling with 800 µM 4-TU was less efficient than expected, most likely due to high and therefore toxic incorporation rates of 4-TU under these conditions.

Finally, we combined both strategies, enhancing pyrimidine salvage and blocking *de novo* biosynthesis, in one cell line: a UMPS knockout line capable of FCU expression, which further enhanced 4-TU incorporation compared to either strategy alone (Figure 1D). As previously reported, we found that UMPS knockout cells exhibited a growth defect compared to the parental cell line (Figure 1E). However, the growth defect could be partially rescued by FCU expression, possibly because FCU increases the supply of pyrimidines via the salvage pathway. RNA-seq analysis of wild-type and UMPS knockout cells revealed a relatively minor effect on transcript levels following UMPS deletion (Supplementary Figure S1C). Furthermore, RNA polymerase II transcripts remained largely unchanged upon pyrimidine starvation in UMPS knockout cells, allowing analysis of their expression under pyrimidine-reduced conditions (Supplementary Figure S1D). In total, only 66 genes transcribed by RNA polymerase II showed deregulation either in UMPS knockout cells or under pyrimidine- reduced conditions. Interestingly, pyrimidine-reduced conditions led to a downregulation of RNA polymerase I transcribed antigen genes in UMPS knockout cells, which are located in the subtelomeric chromosome ends and were not analyzed in this study. Based on the SLAM-seq and RNA-seq data, we decided to perform longer metabolic labeling experiments (7.5 to 60 minutes) in normal HMI-11 medium and to use the pyrimidine-reduced HMI-11 medium only for very short labeling times (≦ 2 minutes) to minimize secondary effects of pyrimidine starvation.

In conclusion, by deleting UMPS and overexpressing FCU, we have established a bloodstream form cell line that allows efficient metabolic RNA labeling. The weak effect of these genetic alterations on the transcriptome and cell growth makes the cell line well suited for precise gene expression studies in *T. brucei*.

### TT-seq suggests unregulated RNA polymerase II transcription in *T. brucei*

To experimentally test the long-standing assumption that RNA polymerase II-mediated transcription is indeed not regulated in *T. brucei*, we assessed the levels of newly transcribed RNA. At very early time points after transcription, before being affected by differences in processing rates, RNA transcript levels reflect the rate of transcription. Thus, knowledge of newly transcribed RNA levels should shed light on whether or not different PTUs are transcribed at similar rates in *T. brucei*. To measure newly transcribed RNA, we established transient transcriptome sequencing (TT-seq) and determined the population of newly transcribed RNA after 0, 30, 60 and 120 seconds of labeling. TT-seq involves the physical separation of newly synthesized RNA from pre-existing, ‘old’ RNA and is well suited to enrich even small amounts of newly synthesized RNA after short labeling times (Figure 2A)(29, 30). To achieve sufficient labeling efficiency after ultra-short periods, labeling was performed in pyrimidine-reduced HMI-11 medium using high 4-TU concentrations (1,600 µM). To ensure rapid cell harvest and avoid prolonged labeling times, cell harvest was performed using a vacuum-based filter system. With this setup, medium was removed and the cells were placed in liquid nitrogen within approximately 5 seconds. Total mRNA was then isolated from the cells, the incorporated 4-TU was biotinylated, and the newly synthesized and labeled RNA was enriched using streptavidin beads. Both, the enriched labeled RNA and the input RNA were sequenced for each replicate, with the input RNA representing total mRNA levels.

**Figure 2.**
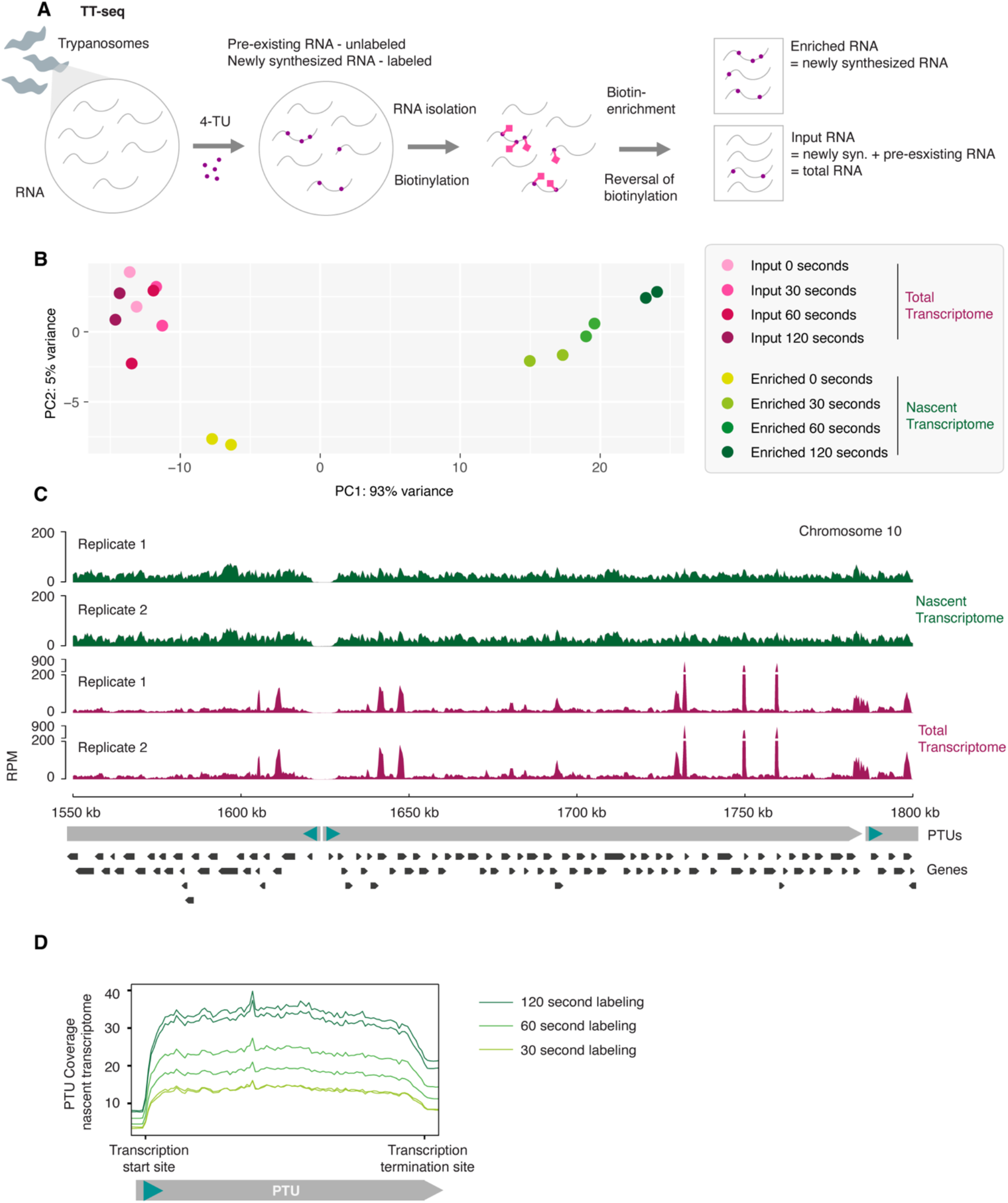
TT-seq suggests that RNA polymerase II transcription is unregulated in *T. brucei*. **(A)** Schematic representation of the transient transcriptome sequencing (TT-seq) approach. TT-seq was performed in UMPS KO cells expressing FCU. Newly synthesized RNA was labeled for 30, 60, and 120 seconds in pyrimidine-reduced medium. The experiment was performed in duplicate. **(B)** PCA analysis of enriched (nascent transcriptome) and input (total transcriptome) samples. **(C)** Visualization of the nascent transcriptome (120 sec labeled, enriched RNA) and the respective total transcriptome (120 sec labeled, input RNA) for a representative core region on chromosome 10. Reads were normalized to reads per million reads mapped (RPM). Both replicates are shown individually. **(D)** Metaplot analysis of nascent transcriptome data along RNA polymerase II transcribed PTUs.

Quality assessment of our TT-seq data by principal component analysis (PCA) revealed marked differences between enriched and input RNA, indicating efficient labeling and specific precipitation of newly synthesized RNA (Figure 2B). A high degree of similarity between the enriched and input samples would have indicated inefficient labeling and unspecific precipitation of the total transcriptome. Consistent with this, the precipitation of unlabeled RNA (0 sec labeling) clustered with the input samples. To support the PCA analysis, we mapped and compared enriched and input transcript levels across the genome (Figure 2C). As expected and previously described (27) for total mRNA levels, we found input transcript levels to vary widely among different genes. In comparison to total mRNA, levels of newly synthesized transcripts were remarkably even among genes (Figure 2C). Taken together, these results indicate that our TT-seq assay had successfully separated newly transcribed RNA from pre-existing RNA, and that our data is well suited to assess nascent RNA transcript levels.

Importantly, beyond serving as a quality control, the finding that newly synthesized transcripts showed almost identical levels across different genes and different PTUs (Figure 2C) provides direct evidence for the long-standing hypothesis of unregulated RNA polymerase II transcription in *T. brucei*. Further, to determine whether transcription elongation was constant along PTUs, we performed metaplot analyses of the nascent transcriptome along all PTUs (Figure 2D). The metaplot analysis indicates that transcription elongation is relatively even along PTUs with no drop in transcript levels towards the end of a unit, as it has been described for RNA polymerase I transcribed subtelomeric PTUs in *T. brucei* (67, 68). Taken together, our TT-seq data support the assumption that RNA polymerase II transcription is unregulated in *T. brucei*, in contrast to most other eukaryotes. Analyzing the nascent transcriptome indicated that RNA polymerase II transcribed PTUs exhibit similar initiation and elongation, resulting in even levels of nascent transcript for each gene.

### RNA stability ranges from several minutes to hours in bloodstream form *T. brucei*

After establishing an efficient metabolic RNA labeling strategy and confirming unregulated RNA polymerase II transcription in *T. brucei* by TT-seq, we set out to measure RNA processing rates and RNA half-lives using the SLAM-seq approach (31, 32). SLAM-seq relies on the chemical conversion of 4-TU labeled RNA rather than its physical separation used in TT-seq. This allows for the detection of both newly synthesized and pre-existing RNA transcripts within one sample (Figure 3A). Metabolic labeling was performed for 0, 7.5, 15, and 60 minutes in standard HMI-11 medium. The time range was chosen because previous RNA half-life measurements suggested a median RNA half-life of 12 minutes in bloodstream form parasites (9). To detect a sufficiently high number of conversion events per sequencing read, we used a high concentration of 4-TU (1,600 µM for 0, 7.5 and 15 min and 800 µM for 60 min labeling) and sequenced the libraries 150 bps from both ends. Old-to-new ratios for RNA polymerase II transcribed genes were calculated in three biological replicates along the time course using the grandR package (44) (Figure 3B). As expected, old-to-new ratios decreased along the time course, reflecting that newly synthesized RNA was produced and replaced pre-existing transcripts that were degraded over time.

**Figure 3.**
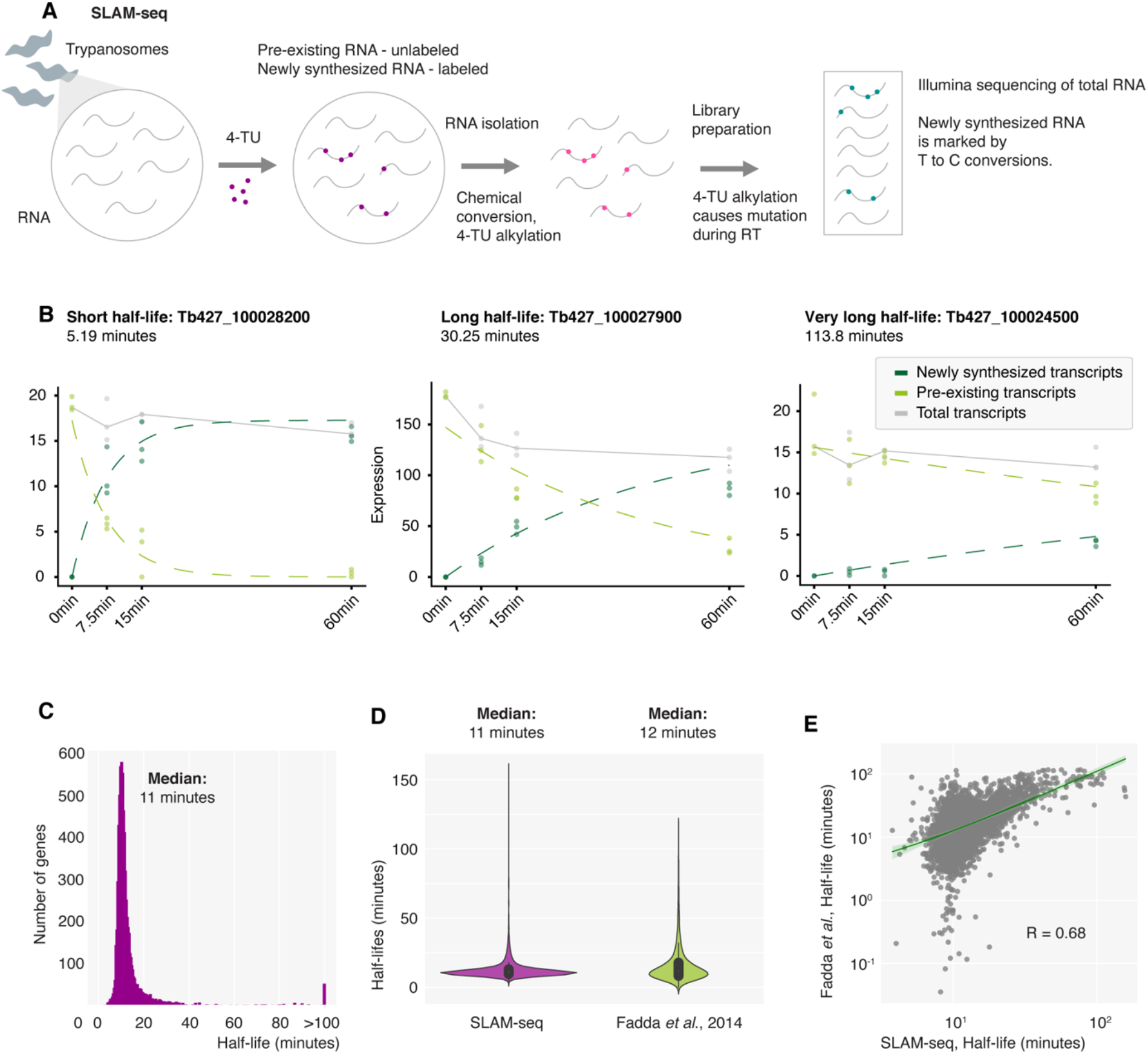
SLAM-seq analysis of RNA half-lives. **(A)** Schematic representation of the thiol (SH)-linked alkylation for metabolic RNA sequencing (SLAM-seq) approach. SLAM-seq was performed in UMPS KO cells expressing FCU using standard HMI-11 medium. Newly synthesized RNA was labeled for 0, 7.5, 15, and 60 minutes in triplicate. (**B**) Old and new RNA were determined by the grandR pipeline based on T-to-C conversion frequency (44). Old, new and total mRNA levels are plotted for three representative genes at different time points. **(C)** Distribution of RNA half-lives measured by SLAM-seq in bloodstream form parasites. **(D)** Violin plot showing RNA half-lives determined by SLAM-seq in this study with RNA half-lives previously determined using a transcription inhibitor and conventional RNA-seq (9). **(E)** Correlation analysis between RNA half-lives determined in this study and RNA half- lives previously determined using a transcription inhibitor and conventional RNA-seq (9).

From the old-to-new ratios, we were able to reliably infer RNA half-lives for most RNA polymerase II transcribed genes (see Supplementary Table S1). In good agreement with the previously published dataset (9), our RNA half-live measurements ranged from a few minutes to several hours, with most half-lives ranging from 3 to 20 minutes (Figure 3C). The median half-life determined by the SLAM-seq experiment was 11 minutes, which is similar to the median of 12 minutes reported by Fadda *et al*. in 2014 (Figure 3D). Fadda *et al.* had used a transcriptional inhibitor followed by standard RNA-seq analyses to infer RNA half-lives. However, while the two datasets compared well globally (R=0.68), the measured rates differed substantially for some genes with very short or very long RNA half-lives (Figure 3E). Many genes with very short half-lives in the previous experiment showed intermediate half-lives in our measurement, suggesting that the extremely short half-lives measured previously may have been a technical artifact. Recently, it has been shown that global inhibition of essential cellular processes, such as transcription, can greatly affect RNA half-lives and therefore interfere with the measurement,(31, 69), for example by accelerating their degradation or stabilizing them by sequestration in biomolecular condensates. Consistent with this, some of the genes that showed a long half-life in the previous experiment showed average stability in our SLAM-seq measurement. In addition, we were able to estimate the half-lives of several very stable genes for which the half-lives have not been previously determined. In the previous study, genes with half-lives exceeding the longest measurement time point of 120 minutes were simply marked as “stable” (9). Since the last time point in the SLAM-seq measurement is 60 minutes, half-lives of >120 minutes could be estimated, although with less precision than shorter RNA half-lives (43).

In summary, using SLAM-seq we were able to successfully measure RNA half-lives in bloodstream form *T. brucei* for 7,314 genes. Compared to a previous measurement (9), secondary effects of global transcriptional inhibition were avoided, allowing for a more accurate measurement.

### RNA processing and RNA stability control total mRNA levels independently

The newly generated SLAM-seq data have not only enabled us to precisely measure RNA half-lives, but also to assess the RNA synthesis rate for each gene, which is influenced by both transcription and RNA processing. Utilizing the grandR pipeline and assuming 20,000 transcripts per cell (9), we estimated the mean RNA synthesis rate for a gene in a single *T. brucei* cell to be about 6.1 transcripts per hour (Figure 4A). Given the parasite’s unregulated transcription that result in even transcription levels for RNA polymerase II genes, variations in mRNA synthesis rates likely reflect the rate of RNA processing, i.e. the rate at which pre- mRNA is converted into mature mRNA through trans-splicing and polyadenylation.

**Figure 4.**
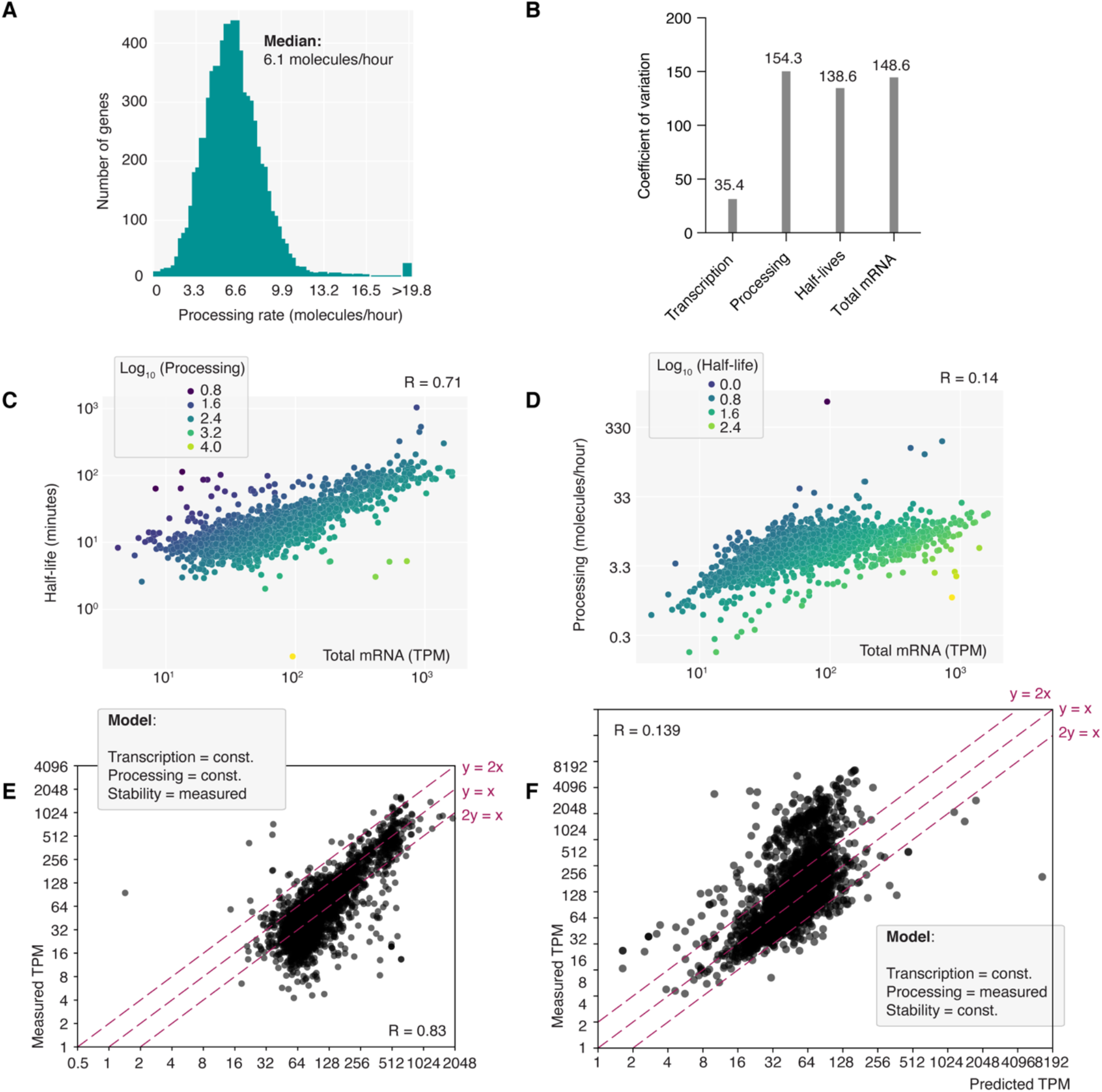
RNA processing and RNA stability can operate independently. **(A)** Distribution of RNA processing rates measured in bloodstream form cells by SLAM-seq. RNA processing rates were inferred from RNA synthesis rates determined by the grandR pipeline (44). **(B)** The coefficient of variation was calculated for transcription (TT-seq data), RNA processing (SLAM-seq data), RNA half-lives (SLAM-seq data) and RNA total levels (SLAM-seq data). (**C**) Correlation analysis of RNA total levels and RNA half-lives, both measured by SLAM-seq. Each dot represents a gene. Dots are colored according to the respective RNA processing rate of the gene. **(D)** Correlation analysis of RNA total levels and RNA processing rates, both measured by SLAM-seq. Each dot represents a gene. Dots are colored according to the respective RNA half-life of the gene. **(E)** Correlation of predicted and measured total mRNA levels. Prediction of total levels was performed assuming constant transcription and constant RNA processing rate. Measured half-lives were introduced in a gene-specific manner. **(F)** Correlation of predicted and measured total mRNA levels. Prediction of total levels was performed assuming constant transcription and constant RNA stability. Measured processing rates were introduced in a gene-specific manner.

Interestingly, our analysis of the coefficient of variation across all genes revealed that RNA processing rates and RNA stability exhibit similar variation levels among individual genes (Figure 4B). In contrast, the variation in gene-specific transcription rates, as determined by TT-seq, was much lower. Since the degree of variation reflects the degree to which a process can influence the output of gene expression, the observed data are consistent with the apparent lack of transcriptional control and suggest that RNA processing and RNA stability influence total mRNA levels to a similar extent. However, correlating these factors with total mRNA levels revealed a much stronger correlation for RNA half-life (R=0.71) than for RNA processing rates (R=0.14) (Figure 4C,D), indicating that differences in RNA half-life more significantly affect RNA levels. Moreover, no correlation was observed between RNA processing rates and RNA stability, suggesting that both processes can occur independently (Supplementary Figure S2A).

To further validate these observations, we employed a computational kinetic model for the *T. brucei* gene expression cascade that has been used in previous studies (9, 57, 59). This model, based on ordinary differential equations, incorporates growth dilution effects, and is parameterized with experimental data. It allows for the simulation of how variations in parameters influence total transcript levels. For this study, we assumed a constant transcription rate across all transcripts, incorporating either measured processing rates or mRNA half-lives, and simulated outcomes for all genes. Consistent with the correlations described above, in our model RNA half-life proved to be a stronger predictor of total mRNA levels than the RNA processing rates (Figure 4E,F). However, none of the two processes was sufficient to fully predict RNA total levels on its own. Instead, both processes exert control, and it is the combination of them that determines RNA total levels.

We also explored potential spatial regulation of the two post-transcriptional processes along the chromosome cores, i.e. the possibility that genomic location affects a gene’s processing or stability (Supplementary Figure S2B). However, when we plotted RNA half-lives and RNA processing rates on the eleven megabase chromosomes of *T. brucei*, we found that the rate of both processes, RNA stability and RNA processing, varied uniformly along the chromosomes. Furthermore, we detected similar median activity for both processes for the different chromosomes. This suggests a lack of spatial regulation for these processes and is consistent with both processes operating in an independent manner.

The lack of correlation between RNA processing rates and RNA half-lives suggests that both processes can operate independently. Therefore, this finding raises the possibility that RNA processing and RNA half-lives may differentially control total mRNA levels for different transcripts.

### RNA processing and RNA stability affect single cell biology

To explore the biological significance of RNA processing and RNA stability occurring independently while both influencing total mRNA levels, we divided genes into four groups based on their RNA processing rates and RNA half-lives relative to the median (Figure 5A). As expected, genes with high processing rates and high RNA stability had the highest total mRNA levels, whereas genes with low processing rates and low RNA stability had the lowest total mRNA levels (Figure 5B). This indicates that RNA processing and RNA stability can synergistically affect RNA levels to produce either very high or very low total mRNA levels. In addition, genes with high processing rates and low RNA stability and genes with low processing rates and high stability showed very similar total mRNA levels, highlighting the fact that there are two alternative pathways for a cell to obtain similar, intermediate total mRNA levels.

**Figure 5.**
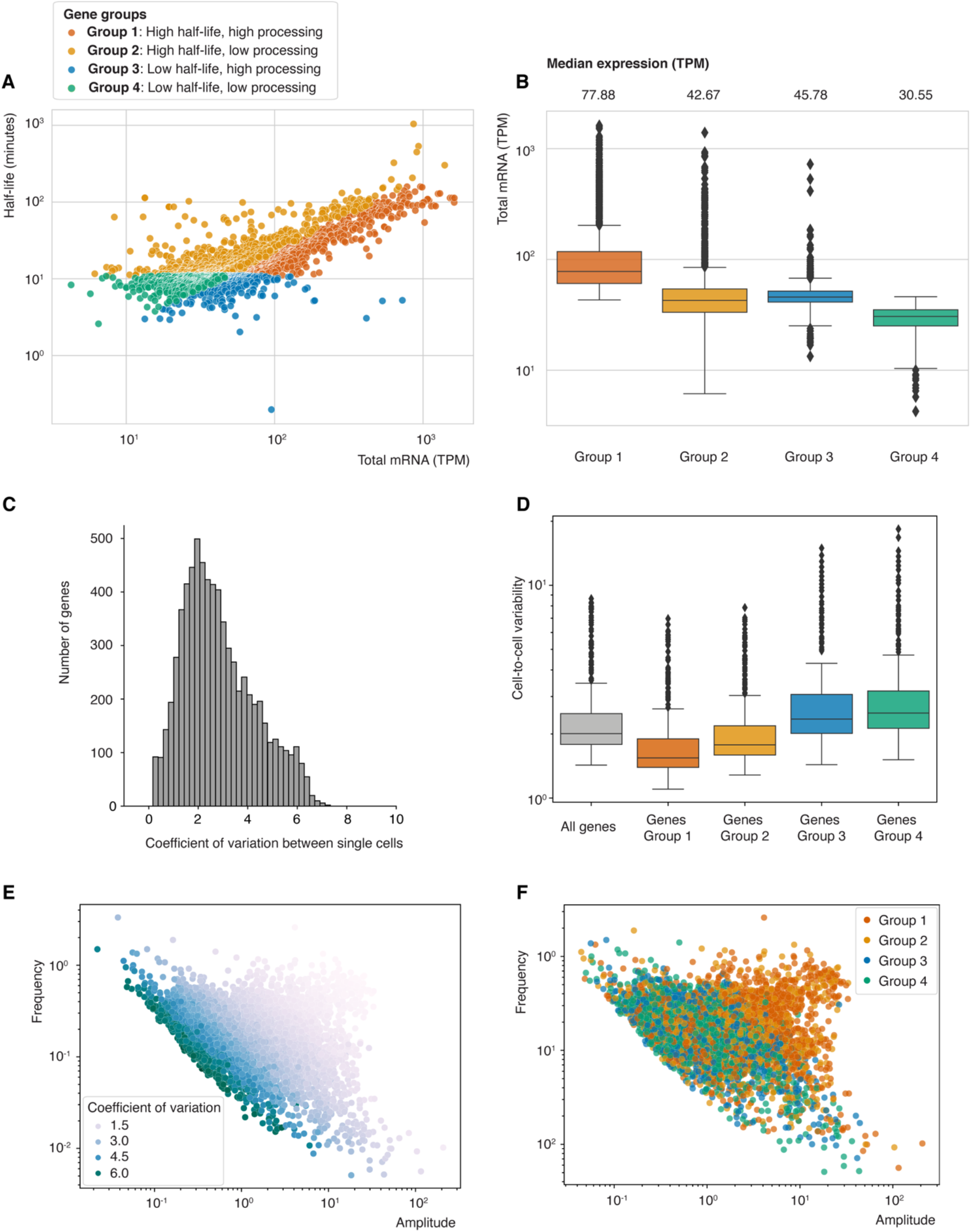
scRNA-seq suggests that RNA processing and RNA stability affect cell-to-cell variability. Genes were classified into four regulatory groups according to their RNA stability and processing rates. Group 1: RNA stability higher than median for all genes, RNA processing rate higher than median for all genes (number of genes = 1,831). Group 2: RNA stability higher than median for all genes, RNA processing rate lower than median for all genes (number of genes = 1,310). Group 3: RNA stability lower than median for all genes, RNA processing rate higher than median for all genes (number of genes = 1,346). Group 4: RNA stability lower than median for all genes, RNA processing rate lower than median for all genes (number of genes = 1,689). **(A)** Visualization of four gene groups. Total levels and RNA stability was correlated for all genes. Each dot represents a gene and is colored according to the group it belongs to. **(B)** Total mRNA levels for each gene group are plotted as box plots. The box represents 25-75 % of the data points, the bar represents the median for the underlying data, the whiskers represent 1.5* interquartile range (IQR) and the rhombi represent outliers. The median total level for each group is shown on top of the graph. **(C)** The coefficient of variation was calculated for each gene in individual cells, using scRNA-seq data from > 300 bloodstream form cells(*60*). The distribution of variation is plotted. **(D)** The coefficients of variation between individual cells are shown as box plots. The box represents 25-75% of the data points, the bar represents the median for the underlying data, the whiskers represent 1.5* IQR and the rhombi represent outliers. The coefficient of variation between individual cells was calculated based on all genes, as well as genes from group 1-4. **(E)** The average detection frequency and amplitude of a gene in individual cells is plotted. Each dot represents a gene and was colored according to coefficient of variation of the gene. The average detection frequency and amplitude of a gene in individual cells was calculated using a generalized telegraph model (GTM) (63). **(F)** The average detection frequency and amplitude of a gene in individual cells is plotted. Each dot represents a gene and was colored according to the regulatory group the gene belongs to.

To understand the ‘benefit’ to a cell of choosing one ‘pathway’ over the other to obtain intermediate transcript levels, we speculated that one path might lead to more robust transcript levels in individual cells than the other, potentially influencing the degree of cell-to-cell heterogeneity. The advent of single-cell RNA-seq (scRNA-seq) has revealed an enormous amount of transcriptional cell-to-cell heterogeneity in biological systems (70, 71). Such cell-to- cell heterogeneity can have advantages and disadvantages for cells and is therefore tightly regulated (70). Because long RNA half-lives increase the time that a transcript is present in individual cells, and therefore increase the chances of detecting a transcript in multiple cells simultaneously, we hypothesized that long half-lives could lead to low cell-to-cell variability for a given transcript. On the other hand, high processing could be an alternative path to reach intermediate total mRNA levels, allowing for more cell-to-cell variability of a given transcript.

To test this hypothesis, we used scRNA-seq data recently generated using a highly sensitive, trypanosome-adapted Smart-seq3xpress protocol (60). The high sensitivity of the approach allowed us to detect on average transcripts from ∼3,000 genes per cell. Based on these data, we analyzed transcript variability across 6,176 genes in 369 cells (Figure 5C). Our findings confirmed that genes with high processing and low stability (group 3) led to greater cell-to-cell variability compared to those with opposite traits (group 2) (Figure 5D-F).

When we performed GO term analyses in order to identify the biological processes enriched in the four different gene groups, we found that genes in groups 1 and 2, which show low cell-to-cell heterogeneity, were involved in core biological processes such as gene expression and cellular transport (Supplementary Figure S3). Genes coding for more specialized cellular processes, such as cell-to-cell communication, were enriched in groups 3 and 4.

In summary, by analyzing cell-to-cell variability using scRNA-seq, we found that RNA stability plays an important role in regulating cell-to-cell variability in *T. brucei*. Furthermore, our results suggest that T*. brucei* may leverage high RNA processing rates and lower RNA stability as an alternative strategy to achieve balanced total mRNA levels, allowing for greater variability among individual cells.

## Discussion

Through genetic re-engineering of the pyrimidine metabolism in *T. brucei*, we have successfully established an efficient metabolic RNA labeling approach, allowing high-resolution analysis of gene expression. Our results provide strong support for the long-standing assumption that RNA polymerase II transcription is unregulated in *T. brucei*. In addition, we were able to quantify RNA stability and processing rates across the genome, providing an invaluable resource for exploring gene expression in this parasite. The new data enabled us to investigate the fundamental question of how these post-transcriptional processes collectively control total mRNA levels. Intriguingly, we observed two distinct paths leading to intermediate total levels: one driven by high RNA stability and the other by high RNA processing rates.

To address the implications of this observation, we analyzed previously generated high-resolution single-cell RNA-seq data and found that genes with intermediate expression levels behave differently at the single-cell level. Genes with high RNA stability and low processing rates showed low variability among cells, whereas genes with low RNA stability and high RNA processing rates showed higher cell-to-cell variability. This suggests that efficient RNA processing may provide an alternative means to achieve intermediate mRNA levels, while allowing for expression variability at the level of single cells. Cell-to-cell variability is thought to be an adaptive feature regulated by cell populations (70, 71) that plays a critical role across the tree of life. Examples of biological processes that depend on the regulation of cell-to-cell variability include the division of labor in bacterial populations, cell differentiation in multicellular organisms, and viral infection (70).

In *T. brucei,* we found that genes associated with high RNA stability and low cell-to-cell variation are involved in core cellular functions such as translation and glucose metabolism. It may be beneficial for individual parasites that such essential mRNAs are present in most cells with low variability, ensuring that basic cellular functions are maintained. Conversely, genes characterized by short RNA half-lives and higher cell-to-cell variability are involved in DNA repair and cell signaling, suggesting a potential advantage for parasite populations to exhibit variability in these processes. Interestingly, throughout its life cycle, *T. brucei* encounters phases in which it benefits from cell-to-cell variability, e.g. in the bloodstream of its mammalian host, where some cells continue to divide as long, slender forms and others differentiate into non-replicative, stumpy forms capable of surviving uptake by an insect vector (71).

In general, transcriptional bursting of individually regulated genes is considered to be the main source of cell-to-cell variability in eukaryotes (70, 72). However, the organization of genes in PTUs, combined with the unregulated transcription of PTUs, limits transcriptional bursting as a source of cell-to-cell variability in *T. brucei*. Our data now suggest that cell-to-cell variation in gene expression may also be generated at a post-transcriptional level. Furthermore, our data suggest that by shifting from high RNA processing to high RNA stability, *T. brucei* may be able to modulate the level of cell-to-cell heterogeneity for distinct sets transcripts despite a lack of transcriptional control. We believe that future studies combining metabolic RNA labeling with single-cell RNA-seq (73) will provide an even better understanding of gene expression dynamics in individual parasites. Such analyses will help to reveal temporal patterns of gene expression and can be used to investigate the role of cell-to-cell variability in parasite survival and disease progression.

## Data Availability

High-throughput sequencing data (TT-seq and SLAM-seq) generated for this study have been deposited in the European Nucleotide Archive under the primary accession number PRJEB71063. Data from the SLAM-seq analysis, such as RNA half-lives and synthesis rates in *T. brucei* bloodstream form cells are available as an interactive resource and for download under: http://einstein.virologie.uni-wuerzburg.de:3839/9faa20e61e41ce24dde7a09ed0191527/

## Code Availability

The code for the analysis of the single cell data is available at: https://github.com/PhysiologicalGenomicBMC/tryp_reproducibility_Luzak2024

## Supplementary Data statement

Supplementary Data are available at NAR Online

## Acknowledgements

We thank all members of the Siegel, Ladurner, Meissner and Boshart laboratories for valuable discussions, T. Straub (Bioinformatics Core Facility, BMC) for providing server space, and the Core Unit LAFUGA at the Gene Center Munich for next-generation sequencing. Furthermore, we thank Zhibek Keneskhanova and Raúl Cosentino for providing scRNA-seq data prepublication; Irina Shcherbakova and Gunnar Schotta for providing the plasmids for RNA spike-in preparation; Christine Clayton for valuable discussions; and R. Cosentino for continuous support with the bioinformatics analysis.

*Author contributions:* The experiments were designed by VL and TNS and carried out by VL unless otherwise indicated. Genetic manipulations to improve RNA labeling were introduced by VL, and growth curves were performed by EO. SLAM-seq and TT-seq experiments were performed by VL with support from EO. TT-seq data were analyzed by VL. SLAM-seq data were analyzed by FE using the grandR tool. Modeling analysis was performed by CO under supervision of JRH. Single-cell RNA-seq analysis was performed by AD with support from SHS. Funding was acquired by TNS, SHS, FE and VL. The work was supervised by TNS. The manuscript was written by VL and TNS and edited by all co-authors. This data was visualized and figures were generated by VL.

---

**Supplementary Figure S1.**
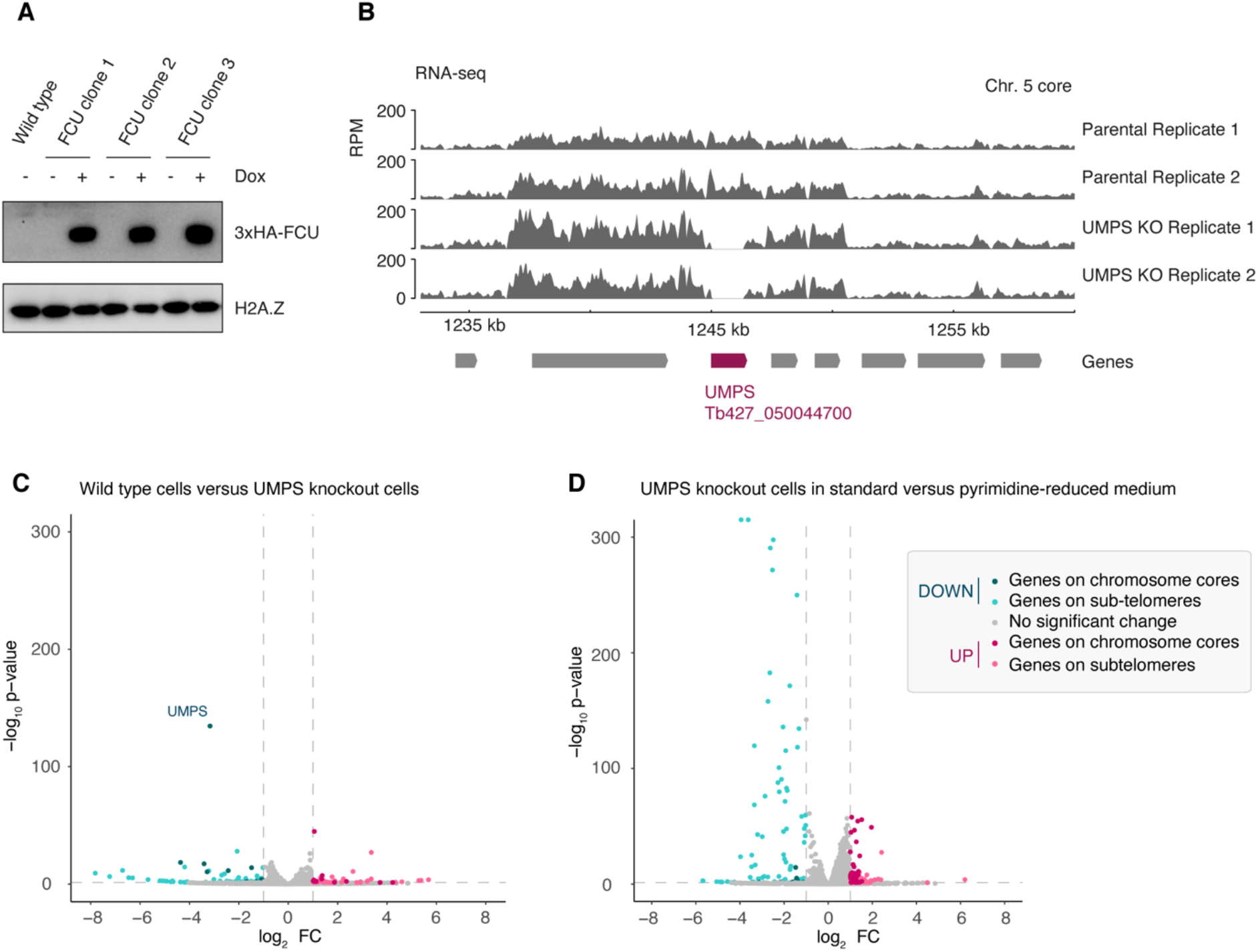
Confirmation of exogenous FCU expression and *UMPS* knockout. **(A)** FCU expression was induced for 24 hours with doxycycline and expression was confirmed by western blot analysis in three independent clones. **(B)** UMPS double knockout was verified by RNA-seq analysis in wild-type and UMPS KO cells. **(C)** Differential gene expression in wild-type versus UMPS KO cells was measured by RNA-seq. The Volcano plot shows unchanged genes in grey, upregulated genes in pink and downregulated genes in green. **(D)** Differential gene expression in UMPS KO cells in standard versus pyrimidine-reduced medium was measured by RNA-seq. Cells were incubated or 15 minutes in either standard or pyrimidine-reduced medium prior to RNA harvest. The Volcano plot shows unchanged genes in grey, upregulated genes in pink and downregulated genes in green.

**Supplementary Figure S2.**
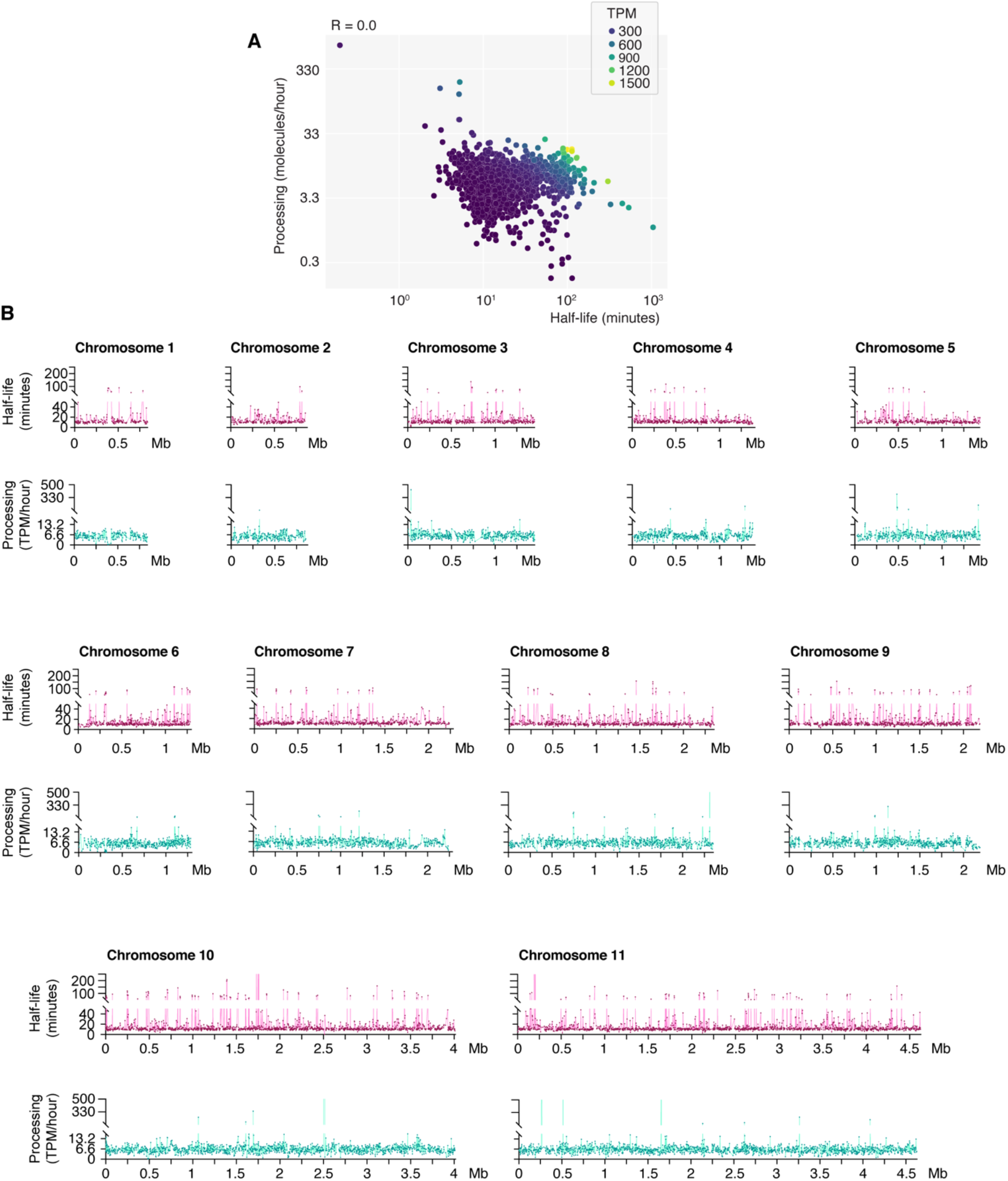
Genome-wide distribution of RNA processing and stability in *T. brucei*. **(A)** Correlation analysis of RNA half-lives and RNA processing rates, both measured by SLAM-seq. Each dot represents a gene. Dots are colored according to the respective RNA total level of the gene. **(B)** RNA processing rates and half-lives were determined for all RNA polymerase II transcribed genes by SLAM-seq, and plotted along the 11 megabase chromosome cores of *T. brucei*.

**Supplementary Figure S3.**
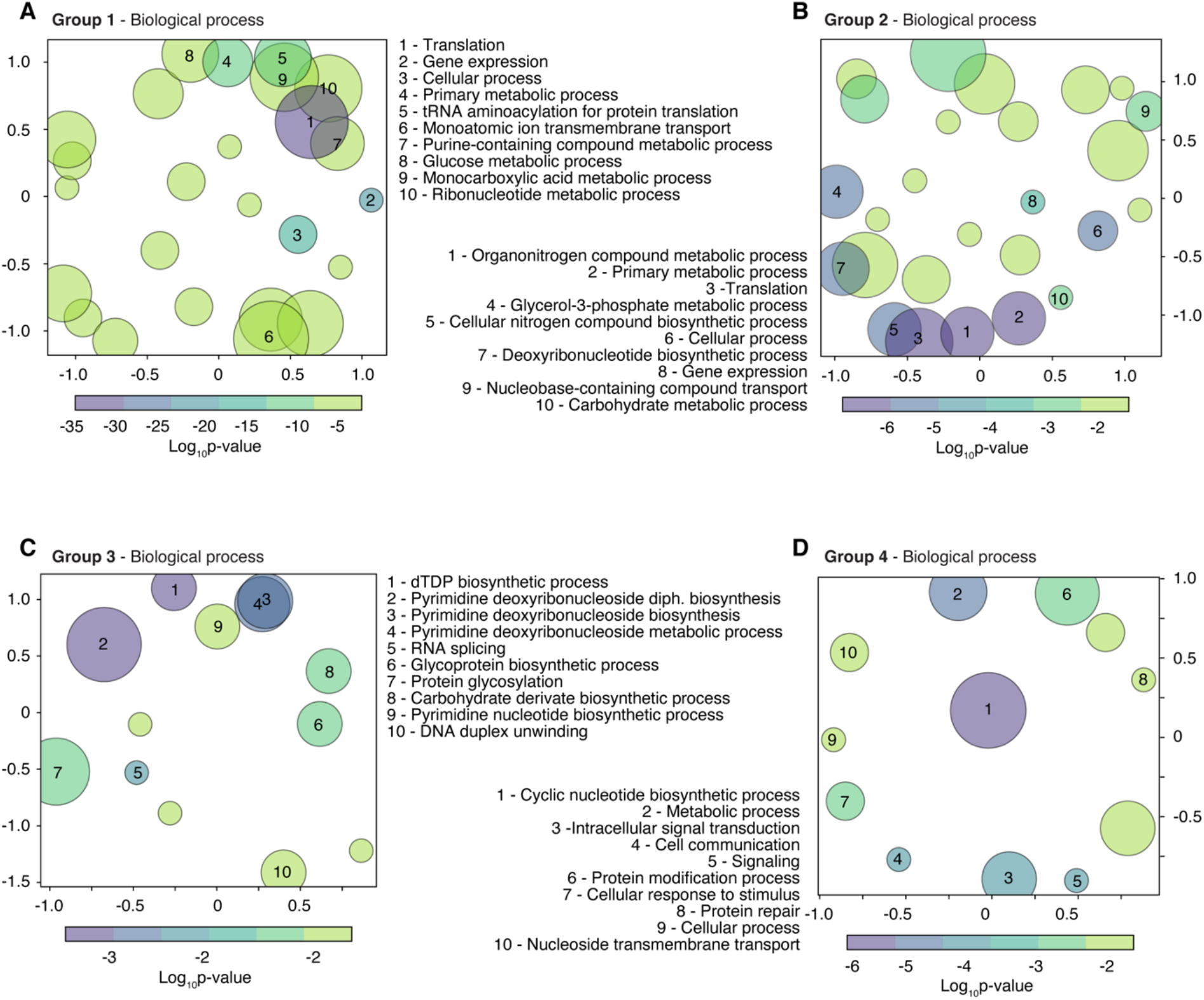
GO-term analysis reveals distinct biological processes represented in four gene groups. Genes were classified into four regulatory groups according to their RNA stability and processing rates. **(A)** GO-term analysis for enrichment of biological processes was performed for 615 genes in group 1 with highest half-lives and highest processing rates. **(B)** GO-term analysis for enrichment of biological processes was performed for 411 genes in group 2 with highest half-lives and lowest processing rates. **(C)** GO-term analysis for enrichment of biological processes was performed for 402 genes in group 3 with lowest half-lives and highest processing rates. **(D)** GO-term analysis for enrichment of biological processes was performed for 323 genes in group 4 with lowest half-lives and lowest processing rates.

## References

1. Orphanides, G. and Reinberg, D. (2002) A Unified Theory of Gene Expression. Cell, 108, 439–451.

2. Picotti, P., Bodenmiller, B., Mueller, L.N., Domon, B. and Aebersold, R. (2009) Full Dynamic Range Proteome Analysis of S. cerevisiae by Targeted Proteomics. Cell, 138, 795– 806.

3. Roeder, R.G. and Rutter, W.J. (1969) Multiple Forms of DNA-dependent RNA Polymerase in Eukaryotic Organisms. Nature, 224, 234–237.

4. Clayton, C.E. (2002) Life without transcriptional control? From fly to man and back again. EMBO J., 21, 1881–1888.

5. Lee, S., Aubee, J.I. and Lai, E.C. (2023) Regulation of alternative splicing and polyadenylation in neurons. Life Sci. Alliance, 6.

6. Bhat, P., Chow, A., Emert, B., Ettlin, O., Quinodoz, S.A., Takei, Y., Huang, W., Blanco, M.R. and Guttman, M. (2023) 3D genome organization around nuclear speckles drives mRNA splicing efficiency. 10.1101/2023.01.04.522632.

7. Matthews, K.R., Tschudi, C. and Ullu, E. (1994) A common pyrimidine-rich motif governs trans-splicing and polyadenylation of tubulin polycistronic pre-mRNA in trypanosomes. Genes Dev., 8, 491–501.

8. Ross, J. (1995) mRNA stability in mammalian cells. Microbiol. Rev., 59, 423–450.

9. Fadda, A., Ryten, M., Droll, D., Rojas, F., Färber, V., Haanstra, J.R., Merce, C., Bakker, B.M., Matthews, K. and Clayton, C. (2014) Transcriptome-wide analysis of trypanosome mRNA decay reveals complex degradation kinetics and suggests a role for co-transcriptional degradation in determining mRNA levels. Mol. Microbiol., 94, 307–326.

10. Ingolia, N.T., Ghaemmaghami, S., Newman, J.R.S. and Weissman, J.S. (2009) Genome-Wide Analysis in Vivo of Translation with Nucleotide Resolution Using Ribosome Profiling. Science, 324, 218–223.

11. Vasquez, J.-J., Hon, C.-C., Vanselow, J.T., Schlosser, A. and Siegel, T.N. (2014) Comparative ribosome profiling reveals extensive translational complexity in different Trypanosoma brucei life cycle stages. Nucleic Acids Res., 42, 3623–3637.

12. Millar, A.H., Heazlewood, J.L., Giglione, C., Holdsworth, M.J., Bachmair, A. and Schulze, W.X. (2019) The Scope, Functions, and Dynamics of Posttranslational Protein Modifications. Annu. Rev. Plant Biol., 70, 119–151.

13. Zhang, N., Jiang, N., Zhang, K., Zheng, L., Zhang, D., Sang, X., Feng, Y., Chen, R., Yang, N., Wang, X., et al. (2020) Landscapes of Protein Posttranslational Modifications of African Trypanosoma Parasites. iScience, 23.

14. Christiano, R., Nagaraj, N., Fröhlich, F. and Walther, T.C. (2014) Global Proteome Turnover Analyses of the Yeasts S. cerevisiae and S. pombe. Cell Rep., 9, 1959–1965.

15. Tinti, M., Güther, M.L.S., Crozier, T.W.M., Lamond, A.I. and Ferguson, M.A.J. (2019) Proteome turnover in the bloodstream and procyclic forms of Trypanosoma brucei measured by quantitative proteomics. Wellcome Open Res., 4, 152.

16. Corchete, L.A., Rojas, E.A., Alonso-López, D., De Las Rivas, J., Gutiérrez, N.C. and Burguillo, F.J. (2020) Systematic comparison and assessment of RNA-seq procedures for gene expression quantitative analysis. Sci. Rep., 10, 19737.

17. Geisel, N. (2011) Constitutive versus Responsive Gene Expression Strategies for Growth in Changing Environments. PLoS ONE, 6, e27033.

18. Lee, T.I. and Young, R.A. (2013) Transcriptional Regulation and Its Misregulation in Disease. Cell, 152, 1237–1251.

19. Gebauer, F., Schwarzl, T., Valcárcel, J. and Hentze, M.W. (2021) RNA-binding proteins in human genetic disease. Nat. Rev. Genet., 22, 185–198.

20. Mignone, F., Gissi, C., Liuni, S. and Pesole, G. (2002) Untranslated regions of mRNAs. Genome Biol., 3, reviews0004.1.

21. Clayton, C. (2019) Regulation of gene expression in trypanosomatids: living with polycistronic transcription. Open Biol., 9, 190072.

22. Presnyak, V., Alhusaini, N., Chen, Y.-H., Martin, S., Morris, N., Kline, N., Olson, S., Weinberg, D., Baker, K.E., Graveley, B.R., et al. (2015) Codon Optimality Is a Major Determinant of mRNA Stability. Cell, 160, 1111–1124.

23. Jeacock, L., Faria, J. and Horn, D. (2018) Codon usage bias controls mRNA and protein abundance in trypanosomatids. eLife, 10.7554/eLife.32496.

24. de Freitas Nascimento, J., Kelly, S., Sunter, J. and Carrington, M. (2018) Codon choice directs constitutive mRNA levels in trypanosomes. eLife, 7, e32467.

25. Cosentino, R.O., Brink, B.G. and Siegel, T.N. (2021) Allele-specific assembly of a eukaryotic genome corrects apparent frameshifts and reveals a lack of nonsense-mediated mRNA decay. NAR Genomics Bioinforma., 3, lqab082.

26. Berriman, M., Ghedin, E., Hertz-Fowler, C., Blandin, G., Renauld, H., Bartholomeu, D.C., Lennard, N.J., Caler, E., Hamlin, N.E., Haas, B., et al. (2005) The Genome of the African Trypanosome Trypanosoma brucei. Science, 309, 416–422.

27. Siegel, T.N., Hekstra, D.R., Wang, X., Dewell, S. and Cross, G.A.M. (2010) Genome-wide analysis of mRNA abundance in two life-cycle stages of Trypanosoma brucei and identification of splicing and polyadenylation sites. Nucleic Acids Res., 38, 4946–4957.

28. Michaeli, S. (2011) Trans-splicing in trypanosomes: machinery and its impact on the parasite transcriptome. Future Microbiol., 6, 459–474.

29. Schwalb, B., Michel, M., Zacher, B., Frühauf, K., Demel, C., Tresch, A., Gagneur, J. and Cramer, P. (2016) TT-seq maps the human transient transcriptome. Science, 352, 1225–1228.

30. Windhager, L., Bonfert, T., Burger, K., Ruzsics, Z., Krebs, S., Kaufmann, S., Malterer, G., L’Hernault, A., Schilhabel, M., Schreiber, S., et al. (2012) Ultrashort and progressive 4sU-tagging reveals key characteristics of RNA processing at nucleotide resolution. Genome Res., 22, 2031–2042.

31. Herzog, V.A., Fasching, N. and Ameres, S.L. (2020) Determining mRNA Stability by Metabolic RNA Labeling and Chemical Nucleoside Conversion. Methods Mol. Biol. Clifton NJ, 2062, 169–189.

32. Herzog, V.A., Reichholf, B., Neumann, T., Rescheneder, P., Bhat, P., Burkard, T.R., Wlotzka, W., von Haeseler, A., Zuber, J. and Ameres, S.L. (2017) Thiol-linked alkylation of RNA to assess expression dynamics. Nat. Methods, 14, 1198–1204.

33. Hirumi, H. and Hirumi, K. (1989) Continuous Cultivation of Trypanosoma brucei Blood Stream Forms in a Medium Containing a Low Concentration of Serum Protein without Feeder Cell Layers. J. Parasitol., 75, 985–989.

34. Scahill, M.D., Pastar, I. and Cross, G.A.M. (2008) CRE recombinase-based positive– negative selection systems for genetic manipulation in Trypanosoma brucei. Mol. Biochem. Parasitol., 157, 73–82.

35. Wirtz, E. and Clayton, C. (1995) Inducible gene expression in trypanosomes mediated by a prokaryotic repressor. Science, 268, 1179–1183.

36. Painter, H.J., Carrasquilla, M. and Llinás, M. (2017) Capturing in vivo RNA transcriptional dynamics from the malaria parasite Plasmodium falciparum. Genome Res., 27, 1074– 1086.

37. Alsford, S. and Horn, D. (2008) Single-locus targeting constructs for reliable regulated RNAi and transgene expression in Trypanosoma brucei. Mol. Biochem. Parasitol., 161, 76– 79.

38. Rico, E., Jeacock, L., Kovářová, J. and Horn, D. (2018) Inducible high-efficiency CRISPR- Cas9-targeted gene editing and precision base editing in African trypanosomes. Sci. Rep., 8, 7960.

39. Wedel, C., Förstner, K.U., Derr, R. and Siegel, T.N. (2017) GT-rich promoters can drive RNA pol II transcription and deposition of H2A.Z in African trypanosomes. EMBO J., 36, 2581–2594.

40. Ali, J.A.M., Creek, D.J., Burgess, K., Allison, H.C., Field, M.C., Mäser, P. and De Koning, H.P. (2013) Pyrimidine salvage in Trypanosoma brucei bloodstream forms and the trypanocidal action of halogenated pyrimidines. Mol. Pharmacol., 83, 439–453.

41. Rädle, B., Rutkowski, A.J., Ruzsics, Z., Friedel, C.C., Koszinowski, U.H. and Dölken, L. (2013) Metabolic labeling of newly transcribed RNA for high resolution gene expression profiling of RNA synthesis, processing and decay in cell culture. J. Vis. Exp. JoVE, 10.3791/50195.

42. Kraus, A.J., Brink, B.G. and Siegel, T.N. (2019) Efficient and specific oligo-based depletion of rRNA. Sci. Rep., 9, 12281.

43. Jürges, C., Dölken, L. and Erhard, F. (2018) Dissecting newly transcribed and old RNA using GRAND-SLAM. Bioinformatics, 34, i218–i226.

44. Rummel, T., Sakellaridi, L. and Erhard, F. (2023) grandR: a comprehensive package for nucleotide conversion RNA-seq data analysis. Nat. Commun., 14, 3559.

45. Dobin, A. and Gingeras, T.R. (2015) Mapping RNA-seq Reads with STAR. Curr. Protoc. Bioinforma. Ed. Board Andreas Baxevanis Al, 51, 11.14.1–11.14.19.

46. Erhard, F., Halenius, A., Zimmermann, C., L’Hernault, A., Kowalewski, D.J., Weekes, M.P., Stevanovic, S., Zimmer, R. and Dölken, L. (2018) Improved Ribo-seq enables identification of cryptic translation events. Nat. Methods, 15, 363–366.

47. Aslett, M., Aurrecoechea, C., Berriman, M., Brestelli, J., Brunk, B.P., Carrington, M., Depledge, D.P., Fischer, S., Gajria, B., Gao, X., et al. (2010) TriTrypDB: a functional genomic resource for the Trypanosomatidae. Nucleic Acids Res., 38, D457–D462.

48. Reijnders, M.J.M.F. and Waterhouse, R.M. (2021) Summary Visualizations of Gene Ontology Terms With GO-Figure! *Front*. Bioinforma., 1, 638255.

49. Ullu, E., Matthews, K.R. and Tschudi, C. (1993) Temporal Order of RNA-Processing Reactions in Trypanosomes: Rapid trans Splicing Precedes Polyadenylation of Newly Synthesized Tubulin Transcripts. Mol. Cell. Biol., 13, 720–725.

50. Gressel, S., Lidschreiber, K. and Cramer, P. (2019) Transient transcriptome sequencing: experimental protocol to monitor genome-wide RNA synthesis including enhancer transcription.

51. Li, H. and Durbin, R. (2009) Fast and accurate short read alignment with Burrows–Wheeler transform. Bioinformatics, 25, 1754–1760.

52. Li, H., Handsaker, B., Wysoker, A., Fennell, T., Ruan, J., Homer, N., Marth, G., Abecasis, G., Durbin, R., and 1000 Genome Project Data Processing Subgroup (2009) The Sequence Alignment/Map format and SAMtools. Bioinforma. Oxf. Engl., 25, 2078–2079.

53. Ramírez, F., Dündar, F., Diehl, S., Grüning, B.A. and Manke, T. (2014) deepTools: a flexible platform for exploring deep-sequencing data. Nucleic Acids Res., 42, W187–W191.

54. Lopez-Delisle, L., Rabbani, L., Wolff, J., Bhardwaj, V., Backofen, R., Grüning, B., Ramírez, F. and Manke, T. (2021) pyGenomeTracks: reproducible plots for multivariate genomic datasets. Bioinformatics, 37, 422–423.

55. Liao, Y., Smyth, G.K. and Shi, W. (2014) featureCounts: an efficient general purpose program for assigning sequence reads to genomic features. Bioinforma. Oxf. Engl., 30, 923–930.

56. Love, M.I., Huber, W. and Anders, S. (2014) Moderated estimation of fold change and dispersion for RNA-seq data with DESeq2. Genome Biol., 15, 550.

57. Antwi, E.B., Haanstra, J.R., Ramasamy, G., Jensen, B., Droll, D., Rojas, F., Minia, I., Terrao, M., Mercé, C., Matthews, K., et al. (2016) Integrative analysis of the Trypanosoma brucei gene expression cascade predicts differential regulation of mRNA processing and unusual control of ribosomal protein expression. BMC Genomics, 17, 306.

58. Olivier, B.G., Rohwer, J.M. and Hofmeyr, J.-H.S. (2005) Modelling cellular systems with PySCeS. Bioinformatics, 21, 560–561.

59. Haanstra, J.R., Stewart, M., Luu, V.-D., Tuijl, A. van, Westerhoff, H.V., Clayton, C. and Bakker, B.M. (2008) Control and Regulation of Gene Expression: QUANTITATIVE ANALYSIS OF THE EXPRESSION OF PHOSPHOGLYCERATE KINASE IN BLOODSTREAM FORM TRYPANOSOMA BRUCEI*. J. Biol. Chem., 283, 2495–2507.

60. McWilliam, K.R., Keneskhanova, Z., Cosentino, R.O., Dobrynin, A., Smith, J.E., Subota, I., Mugnier, M.R., Colomé-Tatché, M. and Siegel, T.N. (2024) High-resolution scRNA-seq reveals genomic determinants of antigen expression hierarchy in African Trypanosomes. 10.1101/2024.03.22.586247.

61. Parekh, S., Ziegenhain, C., Vieth, B., Enard, W. and Hellmann, I. (2018) zUMIs - A fast and flexible pipeline to process RNA sequencing data with UMIs. GigaScience, 7, giy059.

62. Wolf, F.A., Angerer, P. and Theis, F.J. (2018) SCANPY: large-scale single-cell gene expression data analysis. Genome Biol., 19, 15.

63. Luo, S., Zhang, Z., Wang, Z., Yang, X., Chen, X., Zhou, T. and Zhang, J. (2023) Inferring transcriptional bursting kinetics from single-cell snapshot data using a generalized telegraph model. R. Soc. Open Sci., 10, 221057.

64. Pozzi, B., Naguleswaran, A., Florini, F., Rezaei, Z. and Roditi, I. (2023) The RNA export factor TbMex67 connects transcription and RNA export in Trypanosoma brucei and sets boundaries for RNA polymerase I. Nucleic Acids Res., 51, 5177–5192.

65. Pena, A. Role of histone H1 in chromatin and gene expression in the African trypanosome: broad skills, specific functions? (Universidade de Lisboa, 2015)

66. Ong, H.B., Sienkiewicz, N., Wyllie, S., Patterson, S. and Fairlamb, A.H. (2013) Trypanosoma brucei (UMP synthase null mutants) are avirulent in mice, but recover virulence upon prolonged culture in vitro while retaining pyrimidine auxotrophy. Mol. Microbiol., 90, 443–455.

67. Budzak, J., Jones, R., Tschudi, C., Kolev, N.G. and Rudenko, G. (2022) An assembly of nuclear bodies associates with the active VSG expression site in African trypanosomes. Nat. Commun., 13, 101.

68. Weisert, N., Thein, K., Reis, H. and Janzen, C.J. (2021) Quantification of RNA Polymerase I transcriptional attenuation at the active VSG expression site in Trypanosoma brucei. 10.1101/2021.06.21.449234.

69. Haimovich, G., Medina, D.A., Causse, S.Z., Garber, M., Millán-Zambrano, G., Barkai, O., Chávez, S., Pérez-Ortín, J.E., Darzacq, X. and Choder, M. (2013) Gene Expression Is Circular: Factors for mRNA Degradation Also Foster mRNA Synthesis. Cell, 153, 1000–1011.

70. Balázsi, G., van Oudenaarden, A. and Collins, J.J. (2011) Cellular Decision Making and Biological Noise: From Microbes to Mammals. Cell, 144, 910–925.

71. Luzak, V., López-Escobar, L., Siegel, T.N. and Figueiredo, L.M. (2021) Cell-to-Cell Heterogeneity in Trypanosomes. Annu. Rev. Microbiol., 75, 107–128.

72. Larsson, A.J.M., Johnsson, P., Hagemann-Jensen, M., Hartmanis, L., Faridani, O.R., Reinius, B., Segerstolpe, Å., Rivera, C.M., Ren, B. and Sandberg, R. (2019) Genomic encoding of transcriptional burst kinetics. Nature, 565, 251–254.

73. Erhard, F., Baptista, M.A.P., Krammer, T., Hennig, T., Lange, M., Arampatzi, P., Jürges, C.S., Theis, F.J., Saliba, A.-E. and Dölken, L. (2019) scSLAM-seq reveals core features of transcription dynamics in single cells. Nature, 571, 419–423.

